# Genetically modified pigs are protected from classical swine fever virus

**DOI:** 10.1101/361477

**Authors:** Zicong Xie, Daxin Pang, Hongming Yuan, Huping Jiao, Chao Lu, Kankan Wang, Qiangbing Yang, Mengjing Li, Xue Chen, Tingting Yu, Xinrong Chen, Zhen Dai, Yani Peng, Xiaochun Tang, Zhanjun Li, Tiedong Wang, Huancheng Guo, Li Li, Changchun Tu, Liangxue Lai, Hongsheng Ouyang

## Abstract

Classical swine fever (CSF) caused by classical swine fever virus (CSFV) is among the most detrimental diseases, and leads to significant economic losses in the swine industry. Despite efforts by many government authorities try to stamp out the disease from national pig populations, the disease remains widespread. Here, antiviral small hairpin RNAs (shRNAs) were selected and then inserted at the porcine *ROSA26* (p*ROSA26*) locus via a CRISPR/Cas9-mediated knock-in strategy. Finally, anti-CSFV transgenic (TG) pigs were produced by somatic nuclear transfer (SCNT). Importantly, in vitro and in vivo viral challenge assays demonstrated that these TG pigs could effectively limit the growth of CSFV and reduced CSFV-associated clinical signs and mortality, and the disease resistance was stably transmitted to F1-generation. The use of these TG pigs can improve the well-being of livestock and substantially reduce virus-related economic losses. Additionally, this antiviral approach may provide a reference for future antiviral research.

**Author summary:** Classical swine fever (CSF), caused by classical swine fever virus (CSFV), and is a highly contagious, often fatal porcine disease with significant economic losses. Due to its economic importance to the pig industry, the biology and pathogenesis of CSFV have been investigated extensively. Despite efforts by many government authorities to stamp out the disease from national pig populations, the disease remains widespread in some regions and seems to be waiting for the reintroduction and the next round of disease outbreaks. These highlight the necessity and urgency of developing more effective approaches to eradicate the challenging CSFV. In this study, we successfully produced anti-CSFV transgenic pigs and confirmed that these transgenic pigs could effectively limit the growth of CSFV in vivo and in vitro and that the disease resistance traits in the TG founders can be stably transmitted to their F1-generation offspring. This study suggests that these TG pigs can improve the well-being of livestock and contribute to offer potential benefits over commercial vaccination. The use of these TG pigs can improve the well-being of livestock and substantially reduce CSFV-related economic losses.

## Introduction

Classical swine fever virus (CSFV) belongs to the genus Pestivirus within the family Flaviviridae
[1]. CSFV is an enveloped virus that possesses a single-strand positive-sense 12.3 kb RNA genome, which contains a long open reading frame that encodes a poly-protein of 3898 amino acids (aa) [2]. The co- and post-translational processing of the poly-protein by cellular and viral proteases results in the cleavage of the poly-protein into individual proteins [3], including four structural proteins (C, Erns, E1 and E2) and eight non-structural proteins (Npro, p7, NS2, NS3, NS4A, NS4B, NS5A and NS5B) [4].

Classical swine fever (CSF) has a tremendous impact on animal health and the pig industry. CSFV can be transmitted both horizontally and vertically, and domestic pigs and wild boar are highly susceptible to CSFV infection. CSFV can cause severe leukopenia and immunosuppression, which often leads to secondary enteric or respiratory infections [5]. Congenital infection with CSFV can result in persistently infected animals, which do not develop specific antibodies against the virus [6]. This is probably due to immunotolerance developed during foetal lymphocyte differentiation. Persistently infected animals continuously shed virus and are a potential source of new CSF outbreaks, as well as creating problems in diagnosis[7]. Infections with highly virulent CSFV strains show less age-dependent clinical courses and may result in 100% mortality in all age classes of animals and severe neurological signs within 7 to 10 days [8]. The economic losses caused by an outbreak in the Netherlands in 1997 were as high as 2.3 billion USD, and more than 11 million pigs had to be destroyed. When pigs are infected with CSFV strains that have recently circulated in Europe, they become severely ill, and up to 90% die within 4 weeks after infection [9]. Additionally, infected pork and pork products are dangerous sources for the spread of CSF.

Strategies to control CSFV mainly consist of a systematic prophylactic vaccination policy and a non-vaccination stamping-out policy [10]. In 2016, 22 countries officially reported mandatory vaccination campaigns (OIE WAHIS). Compulsory vaccination is the current policy in China, and vaccination coverage must be over 90% at any time of the year in the swine population [11]. Despite efforts by many government authorities to stamp out the disease from national pig populations, the disease remains widespread in some regions [10,12] and seems to be waiting for the reintroduction and the next round of disease outbreaks. These highlight the necessity and urgency of developing more effective approaches to eradicate the challenging CSFV. An alternative strategy is to develop TG pigs that are genetically resistant to CSFV infection.

The rapid development of genome editing technologies has facilitated studies that explore specific gene functions and the establishment of animal models. Recently, the production of genetically modified animals with viral resistance by versatile CRISPR/Cas9 has been examined by several researchers [13,14]. In livestock, these technologies have contributed to the development of antiviral animals and have provided obvious productivity benefits to producers, as well as welfare to the animals. RNAi is a natural post-transcriptional gene silencing mechanism [15], and since its discovery, RNAi has been regarded by virologists as a promising method for the suppression of viral infection[16-19]. To date, the number of reported RNAi-based studies on CSFV suppression in vitro is large, and these studies suggest that developing shRNA-TG pigs that are resistant to viral challenge may be possible.

Therefore, in this study we aimed to combine CRISPR-Cas9 and RNAi technology to generate TG pigs with knock-in of a defined antiviral shRNA, and then assess the transgene resistance of these pigs to CSFV infection (**S 1 Fig**).

## Results

### Selection of antiviral siRNA and shRNA knock-in cell lines

We first aimed to select small interference RNAs (siRNAs) that could efficiently inhibit CSFV. Ten siRNAs (Table 1) were designed and individually introduced into PK-15 cells by electroporation. At 72 h post-infection, the viral inhibitiory efficiencies of these siRNAs were evaluated by qPCR (Fig. 1A) and indirect immunofluorescence assay (IFA) (**S2 Fig**) [20-22]. Of these siRNAs, siRNA-C3 and siRNA-C6 showed higher antiviral ability. Next, we determined whether CSFV could be inhibited in cells stably expressing siRNA-C3 and siRNA-C6. To this end, these two siRNAs were separately cloned into a miR30 recipient vector [23] and formed shRNA targeting donor (**S3A Fig**). The endogenous p*ROSA26* promoter was utilized to express the antiviral shRNA gene (**S3B Fig**).

**Table 1.**
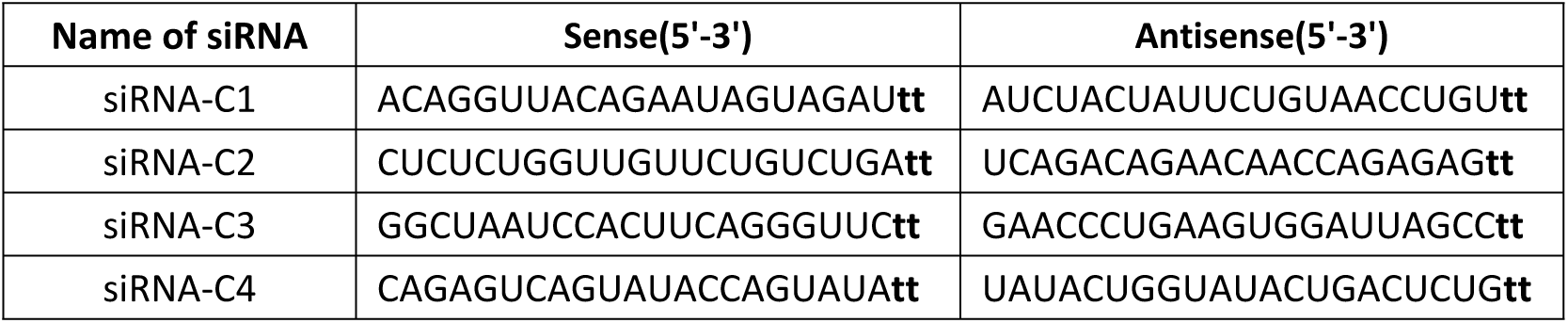
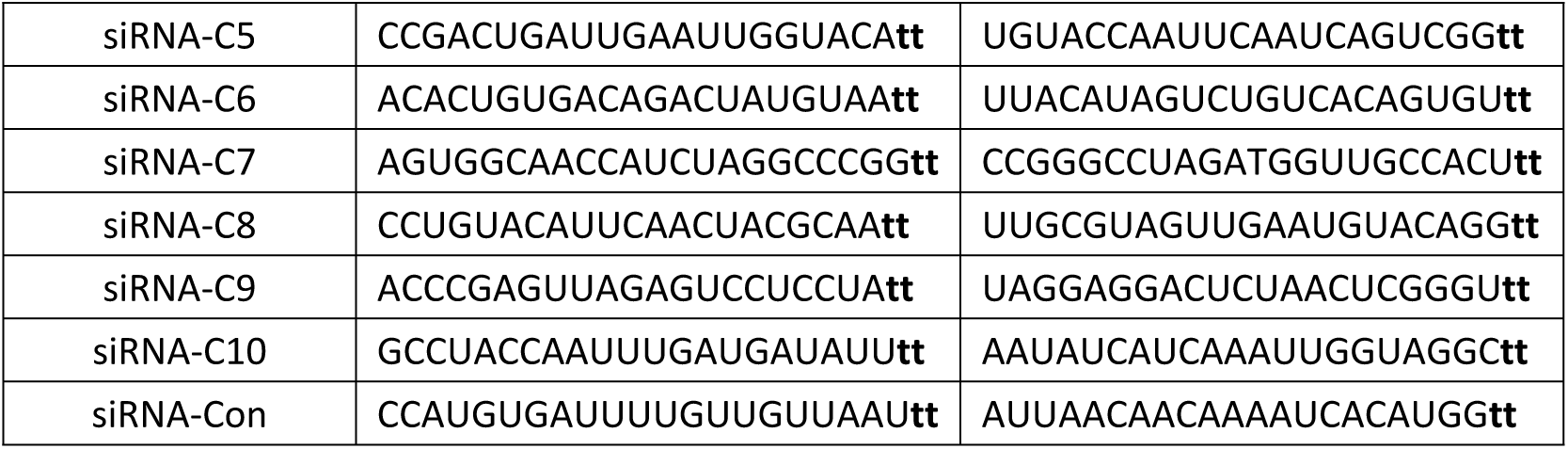
siRNAs sequences.

**Figure 1.**
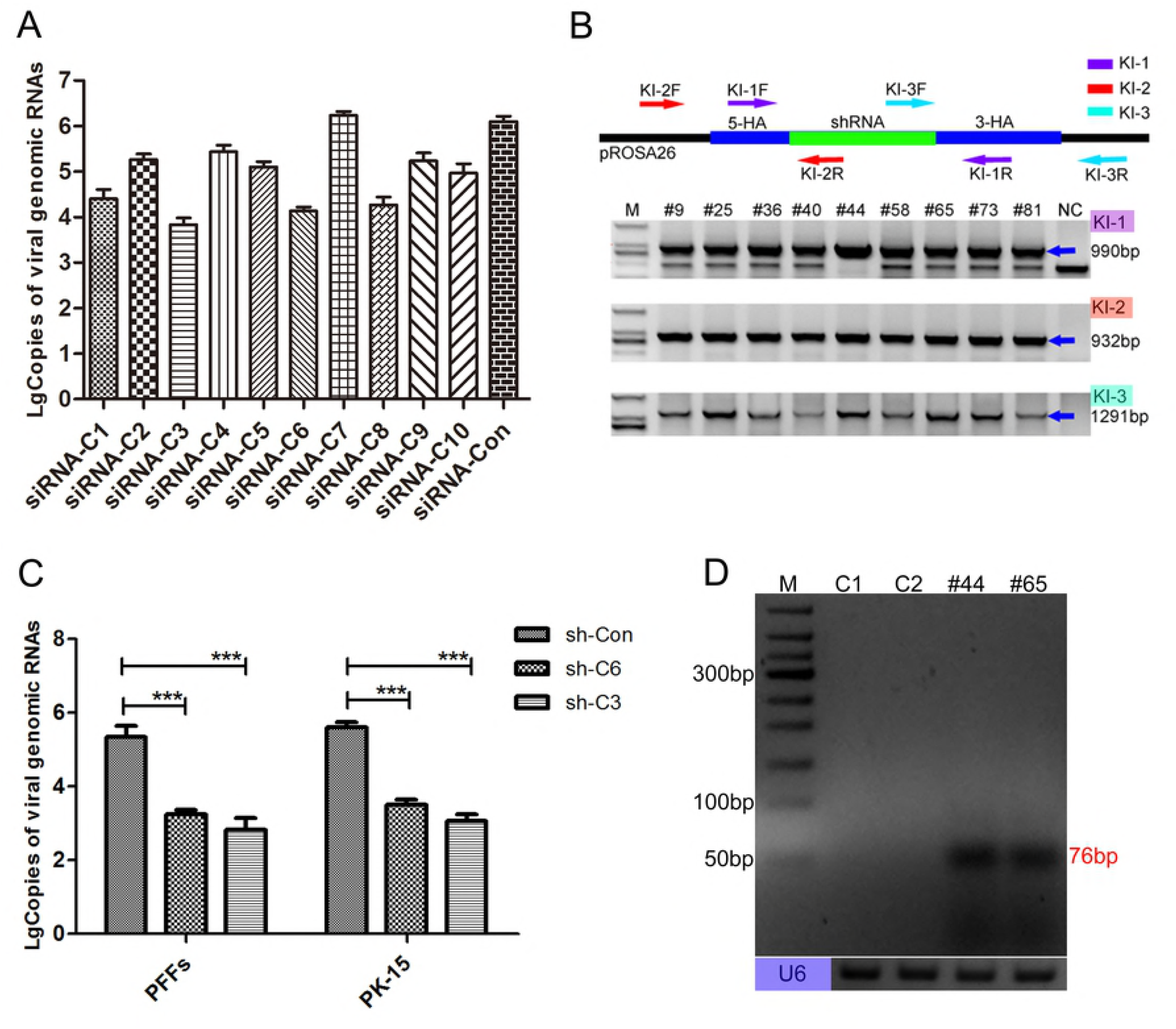
Selection of antiviral cell clones. (**A**) Reduction of viral genome replication in siRNA-transfected cells was further assessed by real time PCR at 72 h post-infection. Error bars represent the SEM, n=3. (**B**) Genomic PCR analysis confirmed the knock-in events at the p*ROSA26* locus by specific primers (**Table 2**). The KI-1 primers were used to determine homozygosity or heterozygosity, the KI-2 primers amplified the 5-junction, and the KI-3 primers amplified the 3’-junction. The blue arrows indicate target amplicons, and the corresponding sizes of the PCR amplicons are 990 bp, 932 bp and 1291 bp, respectively. Lanes 2 – 10 represent the shRNA knock-in positive PFF clones. NC: negative control (wild-type PFFs). M: D2000. The corresponding sequences of these primers are listed in Table 1. (**C**) Inhibition of viable viral production in shRNA knock-in cell clones (PFF cell clones and PK-15 cell clones) were further assessed by real time PCR at 72 h post-infection. The copies of viral genomes were analyzed using the unpaired t-test (^∗∗∗^p<0.001). Error bars represent the SEM, n=3. (**D**) The expression of the two target siRNAs in corresponding transgenic PFF clones was confirmed by RT-PCR. M: 50 bp DNA ladder. C1, wild-type PFFs. C2, scrambled shRNA transgenic PFF clones. #44: shRNA-C3 transgenic PFF clones. #65: shRNA-C6 transgenic PFF clones. The sizes of the target amplicons were 76 bp. Endogenous U6 was used as an RNA quality and loading control.

**Table 2.**
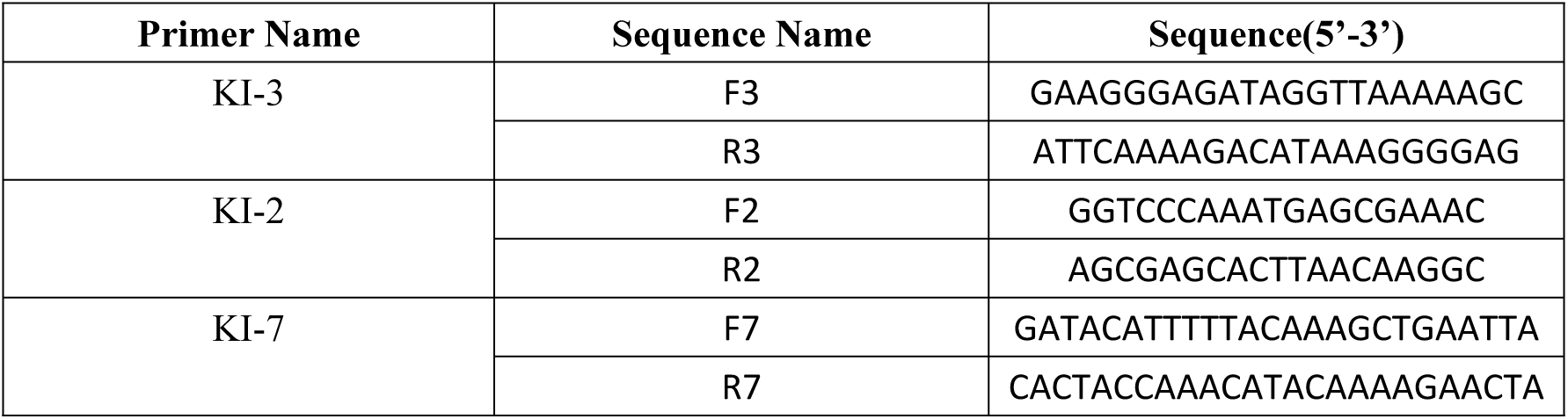
Knock-in primers and corresponding sequences.

Next, these two shRNA targeting donor vectors were separately electroporated into porcine foetal fibroblasts (PFFs) with the CRISPR/Cas9 vector, shRNA knock-in PFFs were selected with the limiting dilution method, and the knock-in events were confirmed by PCR with specific primers and Sanger sequencing analyses (Fig. 1B and **S3C Fig**). Then, we inoculated these knock-in PFFs with CSFV, and 72 h post-infection, the antiviral abilities of these knock-in PFF lines were examined by qPCR (Fig. 1C) and IFA (**S4A Fig**). Similar to the experiments involving transiently transfected siRNA, the viral challenge results indicated that these knock-in PFF lines could effectively inhibit CSFV compared with control PFFs. Finally, the expression of antiviral siRNAs was detected by RT-PCR (Fig. 1D) and further confirmed by sequencing (**S4B Fig**). Additionally, we selected shRNA knock-in PK-15 cells, another porcine cell line that was more susceptible to CSFV infection. As expected, the viral challenge assay of these PK-15 clones showed comparable antiviral ability to that of the PFF clones (Fig. 1C and **S4C Fig**). These results suggested an obvious role for shRNA in inhibiting CSFV growth in these shRNA knock-in cell clones.

To assess potential toxicity caused by shRNA and the further feasibility of generating TG pigs via SCNT, these two shRNA knock-in PFF lines were used as the donor cells for SCNT[24]. The results indicated that the blastocyst development rates were similar between the TG group and wild-type (WT) group (Table 3), suggesting that these knock-in PFFs could be further used to generate anti-CSFV TG pigs via SCNT.

**Table 3.** Statistical results of the blastocyst development rate.

| Repeat Times of SCNT | Type of Donor Cell | Total Nuclear Donor Cell Number | Blastocyst Number | Blastocyst Development Rate |
| --- | --- | --- | --- | --- |
| 1 | TG | 296 | 74 | 25.0% |
|  | WT | 304 | 72 | 23.7% |
| 2 | TG | 311 | 74 | 23.8% |
|  | WT | 306 | 79 | 25.8% |
| 3 | TG | 287 | 70 | 24.4% |
|  | WT | 293 | 73 | 24.9% |

### Production of F0-generation TG pigs and verification of the antiviral ability of isolated TG cells

Then, reconstructed embryos were transferred into surrogate sows, and 8 female TG pigs derived from the same PFF cell clone (shRNA-C3) were successfully obtained from 12 newborns (Fig. 2A). Southern blotting (Fig. 2B) and qPCR (**S6A Fig**) were further designed to investigate whether the shRNA gene was inserted at the pre-determined genome locus and the copy numbers of transgene. As expected, the results confirmed the site-specific integration of antiviral shRNA in the TG pigs. Consistent with the expression pattern of the p*ROSA26* promoter[25,26], we further confirmed the relative expression level of the target siRNA in the various tissues organs and isolated cells from the TG pigs (**S5A Fig**). Next, to investigate whether the isolated TG cells from the TG pigs could be sufficiently effective to inhibit CSFV infection, three different kinds of primary cells were separately isolated from TG pigs and NTG pigs (**S5B Fig**). Then, we inoculated these TG cells with CSFV, and 72 h post-infection, results indicated these isolated TG cells could significantly decrease CSFV compared to NTG cells, as shown in Fig. 2D. Additionally, the replication characteristics of CSFV in the TG and NTG cells were evaluated in a multistep growth curve (Fig. 2E).

**Figure 2.**
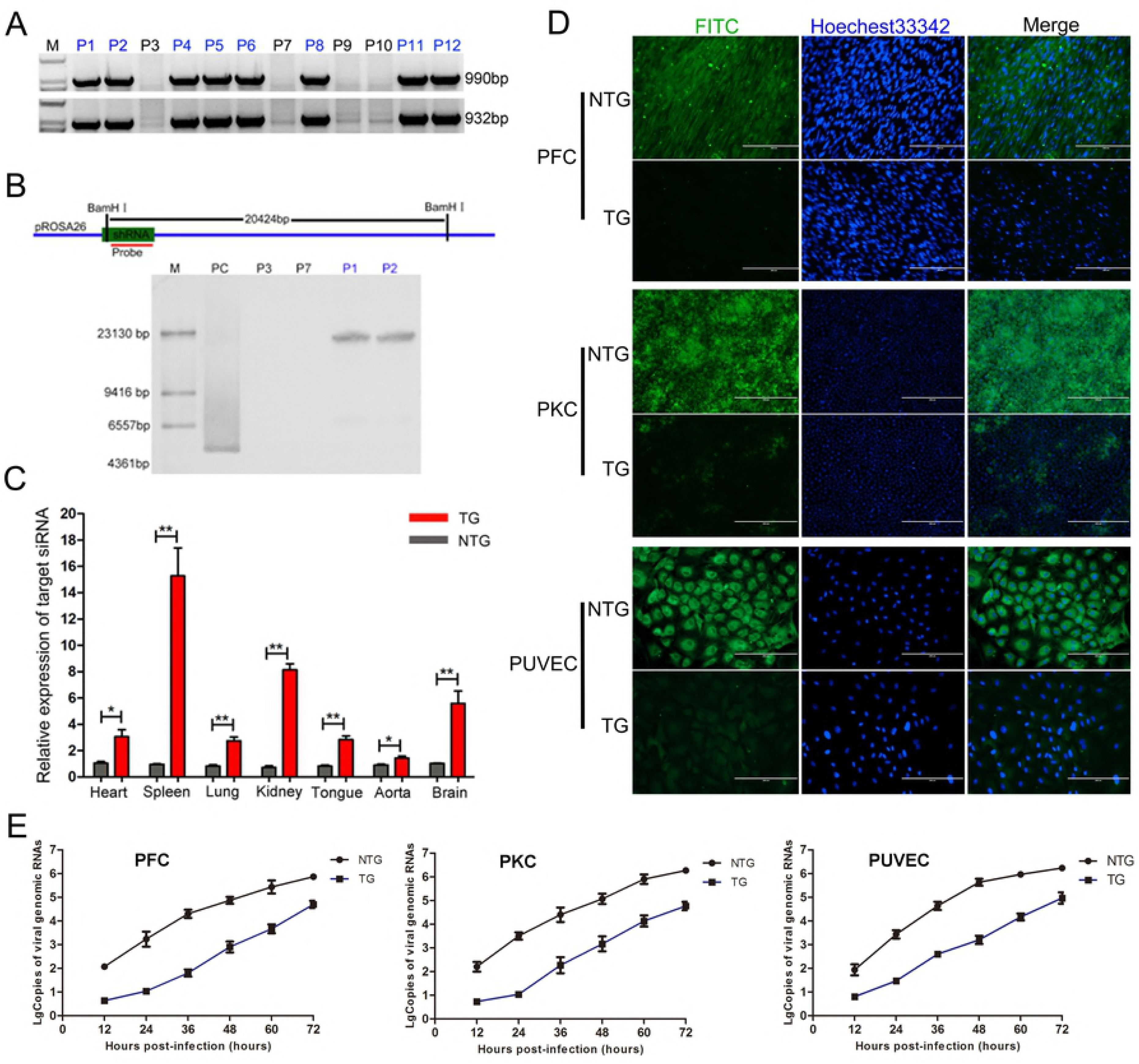
Production of F0-generation TG pigs expressing the target siRNA. (**A**) Genomic PCR analysis of the F0-generation to identify TG pigs. P1, P2, P4, P5, P6, P8, P11 and P12 are TG pigs, and P3, P7, P9 and P10 are NTG pigs. M: D2000. (**B**) Schematic for Southern blotting. The hybridization signals indicated that the shRNA gene was successfully integrated into the p*ROSA26* locus in the TG pigs. M: DNA maker. PC: positive control plasmid. P3 and P7 are NTG pigs, and P1 and P2 are TG pigs. (**C**) The relative expression levels of the target siRNA in various tissues and organs from TG pigs were determined by qPCR. The expression levels were analyzed using the unpaired t-test (^∗^p<0.05; ^∗∗^p<0.01). Error bars represent the SEM, n=3. TG: transgenic pigs. NTG: wild-type pigs. (**D**) Virus resistance in three kinds of isolated primary cells from TG and NTG pigs was examined by IFA. The cells cultured in 24-well plates were inoculated with 200 TCID_50_ of CSFV (shimen strain). IFA was performed using an anti-E2 polyclonal antibody (PAb) and a fluorescein isothiocyanate (FITC)-labeled goat anti-pig IgG (1:100) antibody. PFC: porcine fibroblast cells. PUVEC: porcine umbilical vein endothelial cells. PKC: primary kidney cells. (**E**) Genomic replication kinetics of CSFV in challenged TG and NTG cells at various time points post-infection. Error bars represent the SEM, n=3.

### Generation of TG offspring and analysis of heredity stability

When the TG founders were sexually mature, we obtained 11 F1-generation piglets by crossing the TG founders with WT pigs. Of these piglets, 6 were TG offspring and 5 were NTG offspring (Fig. 3A, B). The TG pigs possessed a stable shRNA gene copy number (**S6A Fig**), a normal porcine diploid chromosome number (2n=38) (**S6B Fig**) and normal birth weight. Additionally, the sex and positive rate of the newborn piglets showed no deviation from the Mendelian law of inheritance. Interestingly, we also found that the molecular beacon could be used to identify the differences in the expression level of the target siRNA between the TG pigs and NTG pigs (**S7 Fig**) and that the expression pattern of the target siRNA in the F1-generation TG pigs was similar to that of their female mother.

**Figure 3.**
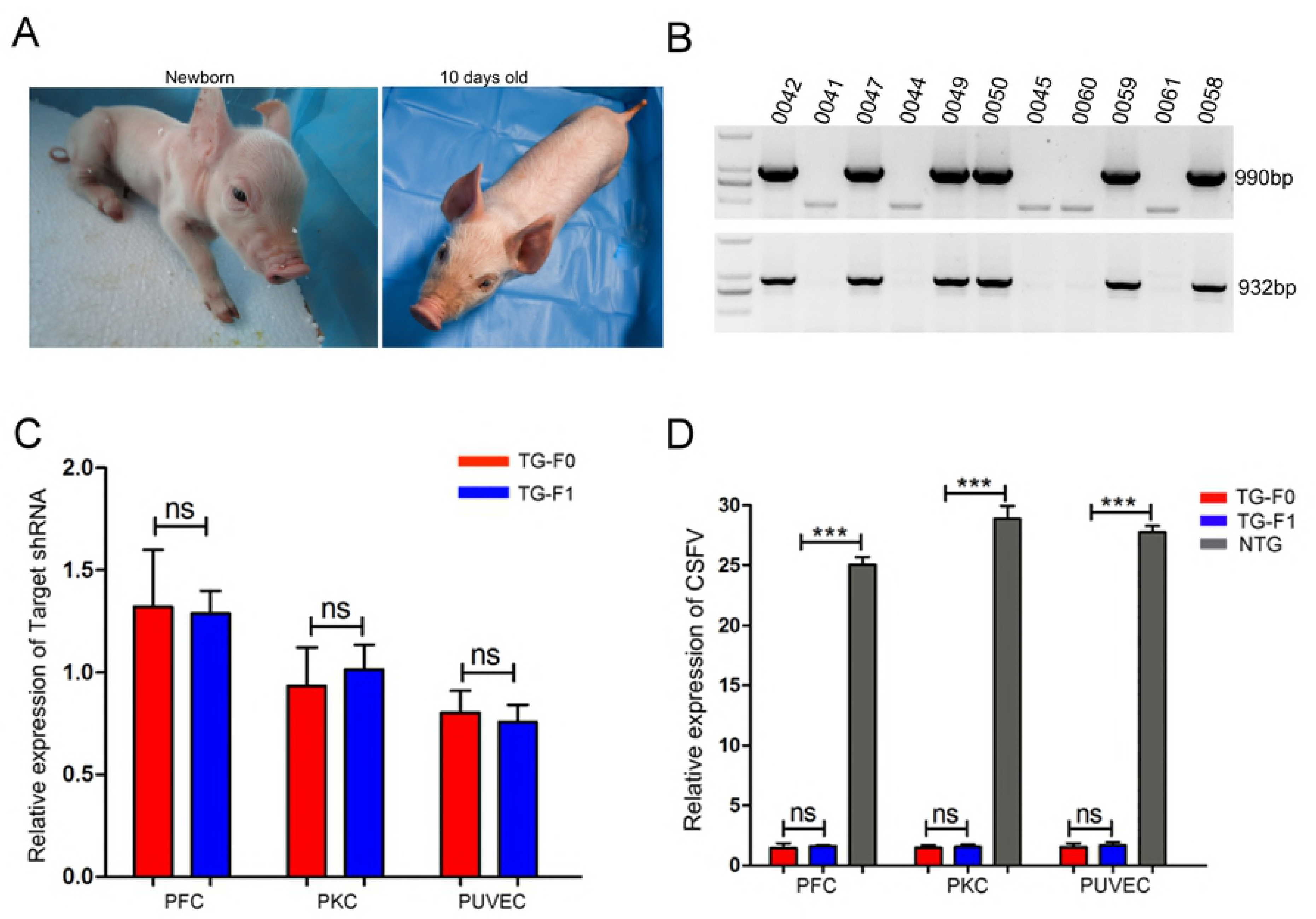
Production of F1-generation TG pigs. (**A**) Photographs show one of the F1-generation TG piglets. (**B**) The knock-in events in newborn TG piglets were confirmed by genomic PCR. Pigs 0042, 0047, 0049, 0050, 0058 and 0059 were TG, and pigs 0041, 0044, 0045, 0060 and 0061 were NTG. M, DNA maker (D2000, TIANGEN, Beijing, China). (**C**) The relative expression levels of the target siRNA in F0- and F1-transgenic cells were compared by qPCR. Target siRNA expression levels were analyzed with the unpaired t-test (ns. not significant). Error bars represent the SEM, n=3. TG-F1 represents F0-generation TG pigs. TG-F1 represents F1-generation TG pigs. (**D**) qPCR results show the relative expression of viral RNA in NTG cells, F0-generation TG cells and F1-generation TG cells. Viral RNA expression was analyzed with an unpaired t-test (ns. not significant; ^∗∗∗^p<0.001). Error bars represent the SEM, n=3.

Importantly, we compared the expression level of the target siRNA between the TG founders and their TG offspring, and the results indicated that there was no significant difference in the shRNA expression level between the F0- and F1-generation TG pigs (Fig. 3C). Moreover, the viral challenge assay showed that the antiviral ability of the parental TG cells was similar to that of their filial TG cells (Fig. 3D, **S6C and S6D Fig**). Together, these findings suggested that the TG pigs could exhibit a higher inhibition to CSFV infection than the NTG pigs and that the RNAi-based antiviral ability could be stably transmitted to their offspring.

### TG pigs resisted CSFV infection

CSF is an acute and highly contagious disease in pigs. In most cases of natural infection, CSFV is mainly transmitted by direct contact with infected animals or by indirect spread via infected feces, food, or carcasses [9]. Therefore, we decided to perform animal challenge experiments in an “in-contact” infectious way. Next, all of the pigs were randomly distributed into two separate rooms, and each room included one non-TG pig that was used to inject CSFV (NTG-In), three NTG pigs and three TG pigs (Fig. 4A). Prior to the viral challenge, all pigs were tested for several common pig-associated viruses (Table 4) and CSFV specific antibody. After three days of acclimation, these NTG-In pigs were directly challenged with CSFV by intramuscular injection, while the other housed pigs were not injected so that the natural spread of CSFV through cohabitation would be mimicked. During the course of infection, CSFV-associated clinical symptoms (including lethargy (**S8A Fig**) and hemorrhage (**S8B Fig**)), mortality and viremia (Fig. 4B) in the challenge test were monitored and recorded daily. Results showed that all of the NTG pigs developed typical signs of CSFV infection, mortality was 100%. Although CSFV-associated clinical symptoms were also observed in the TG pigs, which were not serious, and the virus levels in the blood were lower in the TG pigs than in the NTG pigs.

**Table 4.**
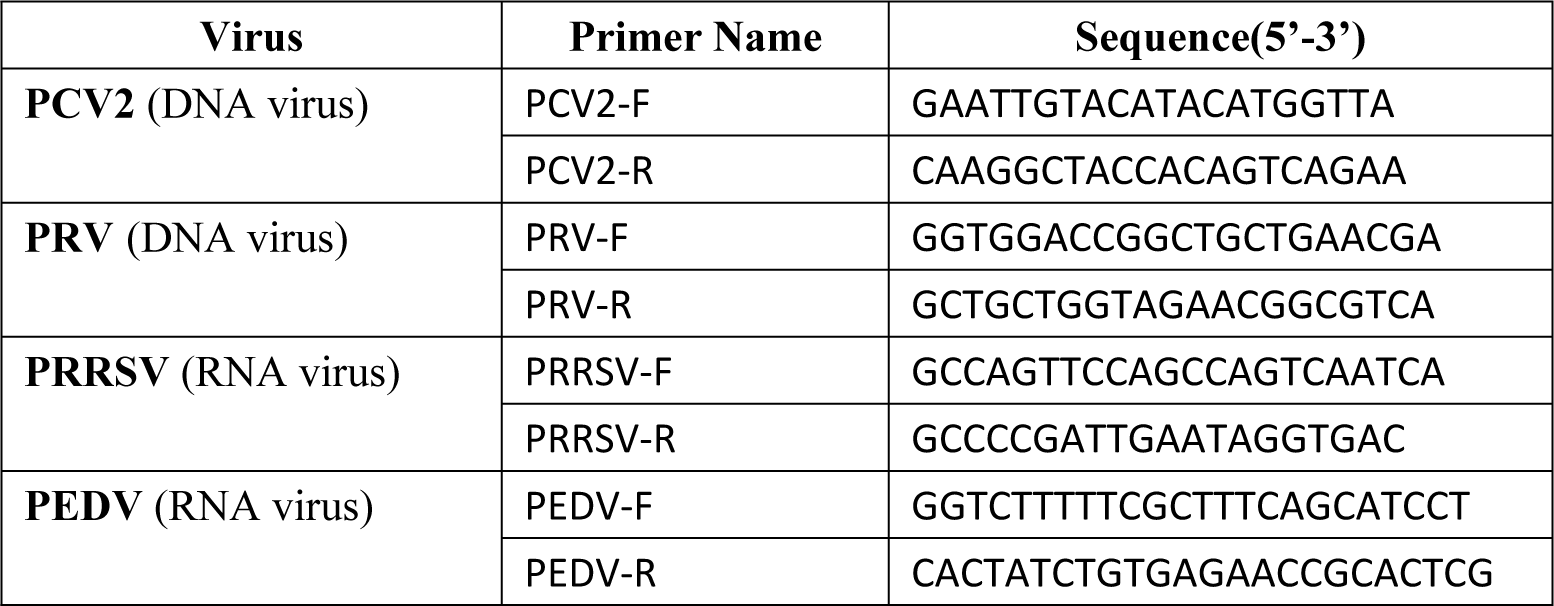
Detection of common viruses in pigs with specific primers.

**Figure 4.**
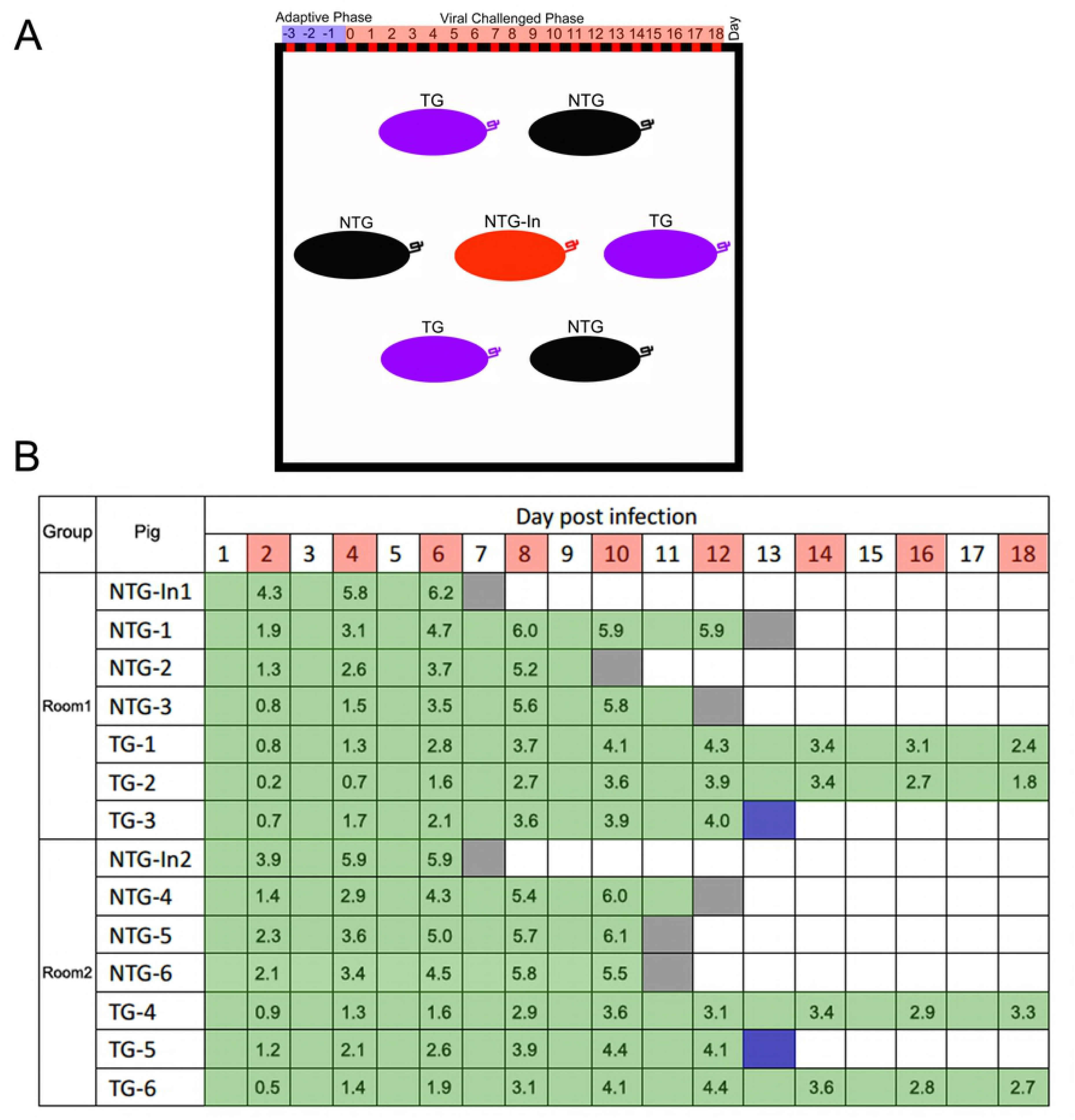
Experimental grouping and the relative clinical monitoring data of the challenged pigs. (**A**) The sketches for the in-contact challenge assay. The black box indicates a separate room. Different colors indicate different types of pigs. Each room included one non-TG pig that was injected with CSFV (NTG-In), three NTG pigs and three TG pigs. (**B**) Mortality and viremia of challenged pigs. The survival time of each pig is indicated by the length of the green bar. Red squares indicate the collection of blood samples. The terminal block color indicates day and cause of death (black, found dead; blue, pigs (TG-4 and TG-5) were euthanized for comparison to the dead NTG pig). Numbers for each day are the lg values of viral RNA copies present in blood samples.

As early as 1 day post-infection (dpi), all of the injected NTG-In pigs developed inappetence, lethargy, and severe diarrhea and fever symptoms. The two injected NTG-In pigs died directly from the infection at 7 dpi, and all of the NTG pigs died between 10 and 13 dpi. By contrast, all of the TG pigs remained alive until euthanasia (Fig. 5A). Additionally, we investigated the body temperatures and daily body weights of these challenged pigs. Compared to the TG pigs, these NTG pigs exhibited more severe fever symptoms (Fig. 5B) and slower weight gain (Fig. 5C). Furthermore, blood samples were collected every 2 days to analyze the dynamic characteristics of the virus in these challenged pigs. Notably, the levels of viral RNA were higher in the NTG pigs than in the TG pigs (Fig. 5D), suggesting that the TG pigs could effectively alter the dynamic characteristics of the viral infection. The antibody response in the challenged pigs was further measured every 4 days until day 16 (Fig. 5E), results showed that the antibody response in the NTG pigs was higher than the TG group. More importantly, compared to the NTG pigs, the antibody response in the TG pigs was delayed by 4~5 days. Meanwhile, compared to the NTG pigs, we found that viral RNA maintained lower levels of expression in CSF susceptible tissues from the TG pigs (Fig. 5F). Additionally, we found that the average times to the initial CSFV-associated clinical manifestation for the NTG and TG were 4 and 8.5 dpi, respectively (**S8C Fig**). Furthermore, when the last two NTG pigs (NTG-1and NTG-3) developed severe symptoms (anorexia, conjunctivitis, severe diarrhea, fever, convulsions and hemorrhage) and were humanely euthanized at 13 dpi, major tissues were collected for histopathological analysis. Finally, the histopathological changes in these tissues (Fig. 5G, **S8D Fig**) were confirmed by hematoxylin/eosin (HE) staining (Fig. 5H). Together, these results indicated that the significant differences in typical signs of CSFV infection and mortality between the TG pigs and NTG pigs during viral challenge, suggesting the antiviral strategy can limit the growth of CSFV in TG pigs.

**Figure 5.**
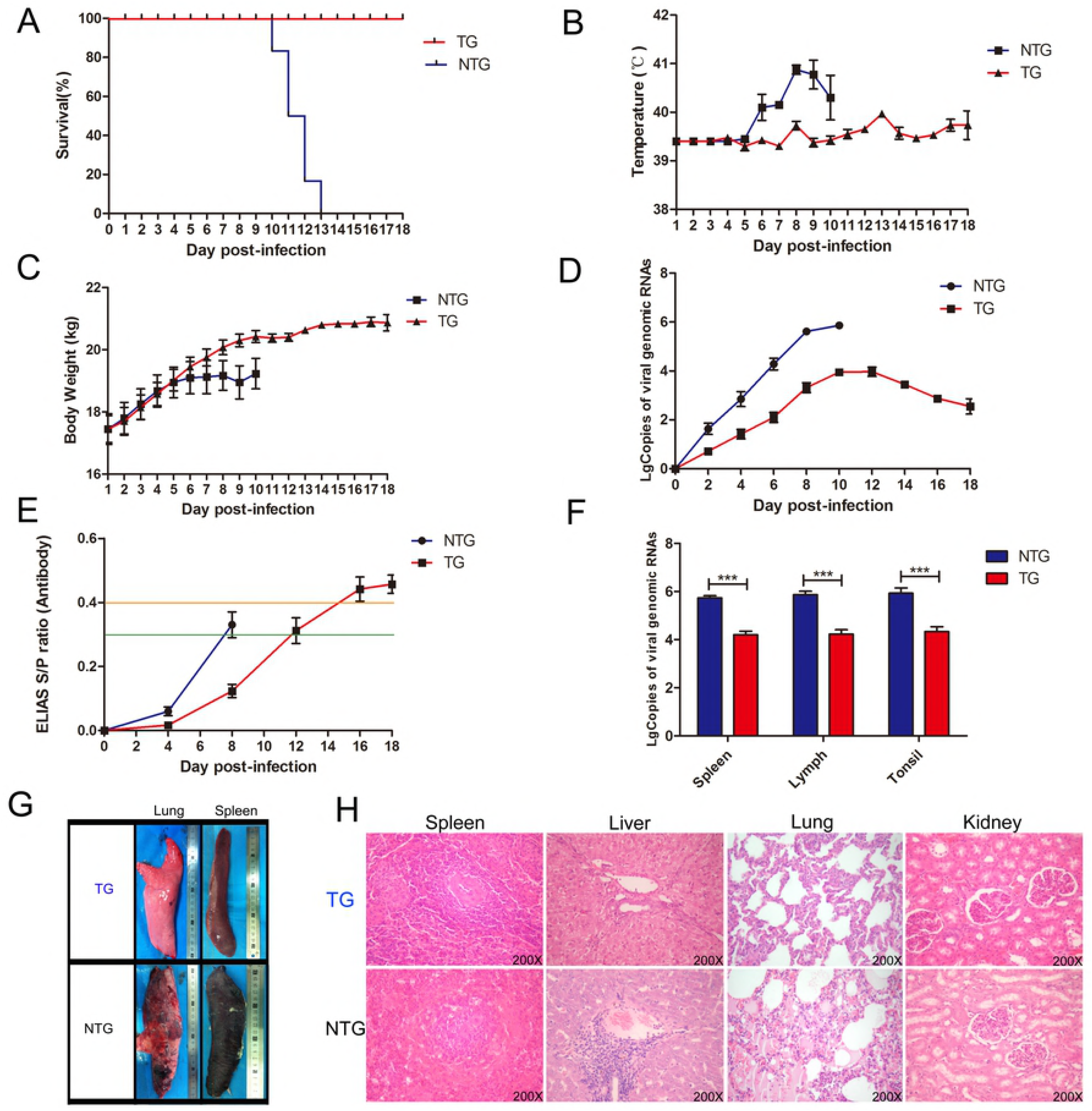
TG pigs exhibit antiviral responses during CSFV infection. (**A**) Survival curves of these challenged pigs during in-contact infection. Different curves indicate different pigs. The blue curve indicates the NTG pigs and the red curve indicates the TG pigs. The pigs in the TG group survived significantly longer than those in NTG group. (**B**) The rectal temperatures of the pigs following challenge with CSFV (TG group, n = 6; NTG group, n = 6). The rectal temperatures were measured daily until the animals died. Error bars represent the standard deviation of at least 3 biological replicates. (**C**) Body weight curves for these challenged pigs during contact infection. Error bars represent the standard deviation of at least 3 biological replicates. (**D**) Viral growth curves in these challenged pigs. CSFV genomic copies in the blood samples are presented according to the standard curve. Data in the panels are presented as the mean ± SE. Error bars represent the standard deviation of at least 3 biological replicates. (**E**) Serum was collected from pigs on 0, 4, 8 and 16 dpi to analyze antibodies against CSFV using a commercial CSFV Antibody Test Kit (IDEXX). ELISA plates were read using a 450 nm filter on an ELISA reader to determine the optical density, and these values were used to calculate the percent inhibition (PI). Error bars represent the standard deviation of at least 3 biological replicates. A sample was considered positive for CSFV antibody when PI was ⩾ 40% (orange Line), Suspicious when was 30% < PI< 40% and negative when PI was optical density ⩽ 30% (green line); (**F**) Viral RNA copies present in major susceptible tissues from the challenged animals (NTG-1 and TG-4) were evaluated by real-time PCR. Viral RNA expression was analyzed with an unpaired t-test (^∗∗∗^p<0.00l). Error bars represent the SEM, n=3. (**G**) Histopathological changes in different tissues and organs between the NTG pigs and TG pigs are indicated by the photos. (**H**) Histopathological changes in the NTG pigs were confirmed by HE staining. These histopathological changes included a decrease in splenic white pulp and hyperemia; the expansion of splenic red pulp in the spleen; acidophilic change and the accumulation of lipid droplets in hepatocytes; the infiltration of inflammatory cells in the portal area of the liver; alveolar effusion, bleeding and the infiltration of a large number of inflammatory cells in the lung; and unclear renal tubular epithelial cell boundaries; and cell cavitation in the kidney.

## Discussion

CSF is one of the most economically important infectious diseases affecting pigs worldwide. Due to its economic importance, intensive control strategies for CSFV have been implemented for several decades, but the disease is still listed by OIE (OIE 2017). There are several possible reasons for the failure to stamp out the disease: (a) the virulence of CSFV is a complex and multifactorial issue that has not been completely defined; (b) the acquired strategies of viral evasion of the host antiviral response still need further in-depth research; (c) the impacts of geography, climate, national policy and people’s awareness of the elimination of disease must be considered; (d) the limitations of current commercial vaccines; (e) the control of wild boar reservoirs is a significant challenge. Additionally, the singleness of control strategies based on vaccination may also be a contributing factor. Therefore, there is still a long way to go toward eliminating the virus. As an alternative, breeding anti-CSFV pigs via a genome editing strategy can be a more direct and effective approach. This would facilitate the permanent introduction of novel disease resistance traits into the mass population of production pigs via conventional breeding techniques.

Recently developed programmable genome editing (PGE) technologies, such as TALEN and CRISPR/Cas9, have been widely used to produce TG animals. Many gene editing strategies can be used to produce viral disease-resistant pigs, and these approaches include knock-out or replacement of attachment factors or receptors (heparan sulfate, sialoadhesin, CD163, etc.) involved in viral infection [27-29], and the inhibition of viral replication via the expression of antiviral gene [30-32]. Compared with other antiviral strategies, RNAi technology has some innate advantages. For example, Regardless of other effects (e.g., the modification of viral receptors may affect normal physiological and biochemical functions, and these modified receptors may be vulnerable to invasion by other diseases), RNAi technology has the flexibility to target one or more loci that are completely conserved and essential for the replication and proliferation of the virus or its serotypes [33]. However, studies on RNAi-mediated anti-CSFV have been mainly focused in vitro, and whether the knock-in of shRNA in pigs can confer permanent resistance against CSFV infection remains unclear[34-36].

In this study, we successfully produced anti-CSFV pigs via the CRISPR/Cas9-based knock-in technology. The in vitro and in vivo viral challenge results demonstrated that these TG pigs exhibited higher resistance to CSFV infection than non-transgenic (NTG) pigs and the acquired RNAi-based antiviral ability in these TG founders could be stably transmitted to their F1-generation offspring. However, even though the extent and severity of clinical signs and viremia were lower in the TG pigs than that in the NTG pigs, some clinic signs and viremia also appeared in the TG pigs. These suggest that the TG pigs combined with other antiviral strategies (systematic prophylactic vaccination, crossing with other type of anti-CSFV pigs, introducing new anti-CSFV features, etc.) can further enhance the antiviral ability. Additionally, we must admit that the animal’s “in-contact” infectious model experiment is small-scale and short-term preliminary studies, because the overall number of TG pigs was low. As encouraging as these results are, a longer-term animal challenge study and the interference with viral transmission in TG pigs need further investigation.

Furthermore, technically, we demonstrated the production of these TG pigs based on CRISPR/Cas9-mediated homology directed repair (HDR). The antiviral shRNA gene in TG pigs was driven by the endogenous p*Rosa26* promoter, which is suitable for driving exogenous gene expression in a consistent and stable manner by avoiding DNA methylation. Additionally, to reduce the potential risk of drug selection and increase the biological safety of the TG pigs, no selectable maker genes were introduced during the production of the TG pigs. This strategy will help provide market support.

Moreover, we believe our antiviral strategy can offers substantial potential benefits over vaccination. The immunization of pigs confers effective protection against CSFV, and the induction of complete clinical protection takes at least 7 days. By that time, the body may become so overrun by infection, and the immune system of the pig may give up trying to battle the infection. In this study, we noticed that the time when the CSFV-associated clinical symptoms began to appear in the TG pigs was significantly delayed (4~5 days) compared with that in the NTG pigs. These finding suggest that these transgenic pigs may have more time to evoke protective immunity and combat the virus. After all, the integrated strategy may be preferred to the singleness of control strategies based on vaccination, and should be considered. Additionally, the CSFV genome is a positive-sense, single-stranded RNA, and it functions as both messenger RNA (mRNA) and a replication template. This TG pig and transgenic strategy can provide some attractive resources to scientists and will help them to better understand and study RNAi.

In summary, we report the combinational application of CRISPR/Cas9 and RNAi in the generation of anti-CSFV TG pigs. We confirmed that these TG pigs could limit the growth of CSFV in vivo and in vitro and that the disease resistance traits in the TG founders can be stably transmitted to their F1-generation offspring. This study suggests that these TG pigs can improve the well-being of livestock and contribute to offer potential benefits over commercial vaccination. The use of these TG pigs can improve the well-being of livestock and substantially reduce CSFV-related economic losses. Additionally, we hope the antiviral approach mentioned in this study will provide some references for future antiviral research.

## Materials and methods

### Cells and virus

Porcine kidney cell line-15 (PK-15) cells (Lot Number: 58808810 ATCC Number: CCL-33^™^) were cultured in Dulbecco’s modified Eagle’s medium (DMEM) supplemented with 5% fetal bovine serum (FBS) (Gibco, Grand Island, New York, USA) and incubated at 39 °C in an atmosphere of 5% CO2. PFF cells were cultured in DMEM supplemented with 10% FBS and incubated at 39 °C in an atmosphere of 5% CO2. PK-15 cells were not included in the list of commonly misidentified cell lines maintained by the International Cell Line Authentication Committee. The origin of the cells (sus scrofa, epithelial) was confirmed by PCR in RIKEN BRC (link of datasheet: http://www2.brc.riken.jp/lab/cell/detail.cgi?cell_no=RCB0534&type=1). The cells were negative for mycoplasma by both PCR and nuclear staining, which were performed based on protocols by RIKEN BRC (http://cell.brc.riken.jp/ja/quality/myco_kensa). CSFV (strain Shimen) and the positive anti-CSFV serum were kindly provided by Dr. Changchun Tu (Academy of Military Medical Sciences, Changchun, China).

### Selection of siRNAs and IFA

All siRNAs were designed and synthesized by Suzhou Genema (Suzhou, China). Then, these siRNAs were individually introduced into PK-15 cells by electroporation, at a siRNA final concentration of 200 nM. Five hours post-transfection, the siRNA-transfected PK-15 cells were inoculated with CSFV and cultured in DMEM with 5% (v/v) fetal bovine serum (FBS) at 39 °C and 5% CO2. 72 hours later, the proliferation of CSFV in siRNA-transfected PK-15 cells was determined by IFA. Briefly, siRNA-transfected PK-15 cells seeded in 24-well plates with four replicates for each siRNA. At 70–80% confluency, the cells were infected with CSFV (200 TCID_50_ per well). At 2 h post-inoculation (hpi), the medium was removed and the cells were cultured in fresh DMEM supplemented with 3% fetal bovine serum. 72 hours later, PK-15 cells were washed three times with cold phosphate-buffered saline (PBS). Then, the cells were fixed in 80% (v/v) cold acetone for at least 30 min in -20 °C/-80°C refrigerator. Next, the fixed cells were washed five times by phosphate-buffered saline with Tween 20 (PBST) and incubated with anti-E2 polyclonal antibody (PAb) (1:100) for 2 h at 37°C, washed five times with PBS, and incubated with a fluorescein isothiocyanate (FITC)-labeled goat anti-pig IgG (1:100) antibody (catalog no. F1638; Sigma-Aldrich) for 30 min at 37°C. After five washes with PBS, the cells were examined using a fluorescence microscope Eclipse TE2000-V (Nikon Imaging, Japan).

### Plasmids

sgRNAs that targeted the p*ROSA26* locus were designed using online software, and sgRNA oligonucleotides were annealed and cloned into the PX330 vector (42230, Addgene) using the method described by Zhang at the Broad Institute of MIT. Targeting sgRNAs were designed and synthesized by Comate Bioscience Co.,Ltd. (Changchun, China). Two complementary sgRNA oligo DNAs were synthesized and then annealed to double-stranded DNA in the presence of 10 × NEB standard Taqbuffer and this product was ligated into the BbsI sites of the vector backbone to form the intact targeting plasmid.

The donor vector contained a 0.5 kb left homology arm (HA) and a 1.0 kb right HA (**Supplemental Figure S3**). The HAs were amplified by genomic PCR and cloned into the PUC57 vector. The shRNA gene was subsequently inserted between the right and left arms.

### Isolation and culture of PFFs

Twelve 33-day-old fetuses were separated from Large White sows in the gestation period, and primary PFFs were isolated from these 33-day-old foetuses of Large White pigs. After removal of the head, tail, limb bones and viscera from the foetal body, the fetuses were cut into small pieces, digested with a sterile collagenase solution and cultured in DMEM supplemented with 20% FBS at 39 °C and 5% CO2 in a humidified incubator.

### Electroporation of PFFs and selection of PFF cell clones

Approximately 3 × 10^6^ PFFs and the corresponding plasmids (30 μg of donor vector, 30 μg of donor vector) were suspended in 300 μL of Opti-MEM (Gibco, Grand Island, New York, USA) in 2 mm gap cuvettes, and electroporated by using specified parameters with a BTX-ECM 2001. The cells were inoculated into ten 100 mm dishes at 48 h post-transfection, and the cell inoculation density per 100 mm dishes was 3,500 cells/dish on average. The cell clones were picked and cultured into 24-well plates. After a confluence of 80% or more was reached, 15% of each cell clone was digested and lysed with 10 μl of NP40 lysis buffer (0.45% NP40plus 0.6% proteinase K) for 1 h at 56 °C and 10 min at 95 °C. The lysate was used as the PCR template and was subjected to 1% agarose gel electrophoresis. Additionally, the knock-in events were confirmed by PCR with specific primers (Table 2). The positive cell clones were thawed and cultured in 12-well plates before SCNT.

### SCNT

The shRNA knock-in positive PFF cells were selected with the limiting dilution method. The positive cells were used for somatic cell nuclear transfer as described previously [24]. Reconstructed embryos were then surgically transferred into the oviducts of surrogate females on the first day of standing estrus. The pregnancy status was monitored using an ultrasound scanner between 30–35 days post-transplantation. Some embryos were cultured for 6-7 days to test the blastocyst formation rate and developmental ability.

### Generation of TG pigs and Southern blotting analyses

shRNA knock-in colonies derived from individual cells were obtained with the limiting dilution method (Xie et al., 2017). These positive cell clones were used as nuclear donor cells to generate transgenic pigs by SCNT. Approximately 250 embryos were transferred into each surrogate pig, and transgenic pigs were delivered by natural birth at full term. Transgene integration was identified by PCR analysis with specific primers. To confirm transgenic insertion into the pig genome, Southern blot was performed by Southern Blot Services (ZooNBIO Biotechnology). DNA was isolated from the TG piglet and WT pig tissues and digested with BamHI. shRNA donor vector was used as a positive control. The probe was hybridized to a 20.424-kb fragment, which is depicted in Figure 2B, indicating site-specific gene insertion.

### siRNA expression level analysis

Small RNAs were isolated by using the miRcute miRNA Isolation Kit (Tiangen, Beijing, China). From purified RNA, complementary DNA was synthesized using the miRcute miRNA First-Strand cDNA Synthesis Kit (Tiangen, Beijing, China). RT-PCR was performed with specific primers. Quantitative RT-PCR was also performed using the miRcute miRNA qPCR Detection Kit (Tiangen, Beijing, China) according the manufacturer’s instructions. SYBR Green real-time PCR was performed using the BIO-RAD IQ^™^5 multicolor real-time PCR detection system. shRNA expression was normalized to the expression of endogenous U6 using the 2–ΔΔCt method.

### Molecular beacon assay

shRNA-specific MB design [37], the MB loop sequence (GGCTAATCCACTTCAGGGTTC) is complementary to the target siRNA, and the MB stem sequence (CCTCC) is typically five nucleotides. Then, an appropriate dye-quencher pair is selected (CY 3 fluorophore & Blank Hole quencher 2), and conjugate the dye and quencher to the 5′ and 3′ ends of the MB sequence, respectively. Prepare total RNA from TG cells and NTG cells, and normalize the total RNA to GAPDH. Then, establish a dose-dependence curve using the serial dilutions of MBs and select optimal concentration for further testing. The MB signal at the highest target oligonucleotide concentration should generally be 5–30 times higher than the background signal quantified in the negative control experiment in which the signal level of MBs without any target is measured. Add 50 μl of MB solution to each well of a 96-well black-bottomed plate, and then add 50 μl solutions containing the target oligonucleotide to their designated wells. Incubate at 37 °C for 5 min to allow the solutions to equilibrate. The fluorescence intensity of MBs is detected by using a microplate reader.

### Determination of transgene copy number

The copy number of antiviral shRNA gene was determined by qPCR as previously described [38]. Briefly, a standard curve was produced with series of standard samples containing 0, 1, 2, 4, 8, 10 copies of the shRNA gene, respectively, by mixing the wild-type genome of pig with shRNA expression vector. The absolute quantitative standard curve was drawn by plotting ÄCt=Ct_shRNA_–Ct_TFRC_ against the log of shRNA gene copies of corresponding standard samples.

### Viral challenge assay in TG cells

The in vitro viral challenge assay was strictly performed at a designated safe place. TG fibroblasts, TG kidney cells and TG umbilical vein endothelial cells were isolated from newborn TG pigs. These cells, cultured in 24-well plates, were inoculated with 200 TCID_50_ of CSFV (Shimen strain), and there were four replicates for each TG cell types. One hour later, the inoculums were replaced with fresh medium (5% fetal bovine serum). After 48-h incubation, cells and virus were collected and evaluated by IFA and qPCR. To analyze CSFV proliferation in TG cells by qPCR, total RNA was extracted from the CSFV-infected cells using TRIzol-A+ reagent (Tiangen, Beijing, China) and reverse transcribed into cDNA using the BioRT cDNA First Strand Synthesis Kit (Bioer, Hangzhou, China) according to the manufacturer’s protocol. SYBR Green real-time PCR was performed using the BIO-RAD IQ^™^5 multicolor real-time PCR detection system and the BioEasy SYBR Green I real-time PCR kit.

### Viral challenge assay in TG pigs

All animal studies were performed according to protocols approved by the animal Welfare Committee of China Agricultural University. All pigs (the NTG-In group (n=2), NTG group (n=6) and TG group (n=6)) were 55 days old and separated into two rooms. The pigs in the NTG and TG groups were same age pigs from the F0 generation of the TG founders. Before the CSFV challenge, all pigs were confirmed to be CSFV negative, and a commercial CSFV enzyme-linked immunosorbent assay kit (ELISA; IDEXX Laboratories, Inc., Westbrook, ME, USA) was used to test CSFV antibodies in these pigs. The NTG-In pigs were challenged by intramuscular injection in the neck with 1.0×10^4^ TCID_50_ CSFV Shimen in 2.5 ml of PBS. The in vivo viral challenge assay was strictly performed at a designated safe place. Then, the transgenic animal corpses were received humane treatment when the experiments were completed.

### Quantification of serum viral RNA

Quantitative RT-PCR was performed to examine CSFV in pig blood. Blood samples from each pig were collected at days 0, 2, 4, 6, 8, 10, 12, 14, 16 and 18 after injection. Viral genomic RNA was isolated by using Trizol (Tiangen, Beijing, China) according to the manufacturer’s instructions. A standard curve was generated to detect the viral load in each blood sample with 10-fold serial dilutions of viral lysates ranging from 10^8^ to 10^2^. SYBR Green real-time PCR was performed using the BIO-RAD IQ^™^5 multicolor real-time PCR detection system and the BioEasy SYBR Green I real-time PCR kit and the Ct values and CSFV RNA copies were determined.

### Histopathological analysis

All animals were killed on the 10th day post-infection. Major tissues, including the heart, spleen, lung and other tissues, from the pigs were fixed in formalin followed by routine paraffin sectioning and HE staining. Histopathological changes were observed under a microscope.

### Ethics statement

All animal studies were approved by the Animal Welfare and Research Ethics Committee at Jilin University (Approval ID: 20160602), and all procedures were conducted strictly in accordance with the Guide for the Care and Use of Laboratory Animals. All surgeries were performed under anesthesia, and every effort was made to minimize animal suffering.

## Supporting Information Legends

**Supplementary Figure 1.** Main technical routes about antiviral TG pig.

**Supplementary Figure 2.** Antiviral ability of various designed siRNAs (siRNA-C1~siRNA-C10) was assessed by IFA in siRNA-transfected PK-15 cells at 72 h post-infection. The cells cultured in 24-well plates were inoculated with 1000 TCID_50_ of CSFV (shimen strain). IFA was performed using an anti-E2 polyclonal antibody (PAb) and a fluorescein isothiocyanate (FITC)-labeled goat anti-pig IgG (1:100) antibody (Sigma). siRNA-Con: scrambled siRNA. NC: negative control (no CSFV).

**Supplementary Figure 3. Verification of site-specific knock-in events for PFF cell clones. (a)** Composition and structure of donor vector for knock-in. 5’HA: left homologous arm; 3’HA: right homologous arm; shRNA: anti-CSFV shRNA gene cassette. (**b**) The scheme for shRNA site-specific knock-in. HA: Homology arm. (**c**) Sanger sequencing analyses were used to further confirm the EGFP site-specific knock-in events in pROSA26 locus.

**Supplementary Figure 4. The expression of target siRNA and verification of antiviral ability in TG PK-15 cell clones.** (a) Virus resistance in shRNA-C3 (#44) and shRNA-C6 (#65) transgenic PFFs was examined by IFA. shRNA-Con: scrambled shRNA transgenic PFFs. (b) Sanger sequencing analyses were used to further confirm the expression of target siRNA in positive PFF cell clones. (c) The replication and proliferation of CSFV in TG PK-15 cell clones were evaluated by IFA. The cells cultured in 24-well plates were inoculated with 200 TCID_50_ of CSFV (shimen strain). IFA was performed using an anti-E2 polyclonal antibody (PAb) and a fluorescein isothiocyanate (FITC)-labeled goat anti-pig IgG (1:100) antibody. shRNA-C3, shRNA-C3 knock-in PK-15 cells. shRNA-C6, shRNA-C6 knock-in PK-15 cells. shRNA-Con, scrambled shRNA knock-in PK-15 cells.

**Supplementary Figure 5. Phenotypic analyses for TG pigs. (a)** Relative expression level of target siRNA (siRNA-C3) in various tissues and cells from TG pigs were detected by RT-PCR. (**b**) Three kinds of primary TG cells isolated from TG pigs. In particular, the isolated PUVECs were labeled with anti-CD31 antibody and performed immunofluorescent analysis.

**Supplementary Figure 6. Phenotypic analyses for F1 generation TG pigs. (a)** The knock-in event of shRNA gene at the pROSA26 locus in F1 generation TG pigs was confirmed by qPCR. Pigs 3900, 3902 and 3904 were F0-generation TG pigs, pigs 0042, 0049 and 0058 were F1-generation TG pigs and pigs 0044 was NTG pigs. Data are means of three replicates±SD. (**b**) Karyotype analysis results indicated that these TG pigs had normal porcine diploid chromosome number (2n=38). (**c**) The viral infection in isolated F1 generation primary TG cells was confirmed by RT-PCR. (**d**) The viral infection in isolated F1 generation primary TG cells was further confirmed by IFA. The cells cultured in 24-well plates were inoculated with 200 TCID_50_ of CSFV (shimen strain). IFA was performed using an anti-E2 polyclonal antibody (PAb) and a fluorescein isothiocyanate (FITC)-labeled goat anti-pig IgG (1:100) antibody.

**Supplementary Figure 7. (a)** Schematic for molecular beacons to detect the target siRNA in TG pigs. (**b**) The relative expression levels of the target siRNA in various tissues and cells in TG pigs and NTG pigs were detected with molecular beacons. TG: transgenic pigs. NTG: wild-type pigs. Target siRNA expression was analyzed with an unpaired t-test (^∗∗^p<0.01; ^∗∗∗^p<0.001). Error bars represent the SEM, n=3.

**Supplementary Figure 8. The results of in vivo viral challenge assay. (a)** Different mental state was observed among these challenged pigs. NTG indicated NTG pigs and TG indicated TG pigs. (**b**) Haemorrhagic signs were observed in different organs and tissues in NTG pigs. ① indicated skin, ② indicated lymph and ③ indicated spleen. (**c**) Statistic data about the time of initial morbidity among challenged pigs. n = 6. Graphs show the mean ± S.E.M. (**d**) Pathological changes were also observed in lymphoid tissue.

